# The role of sensory uncertainty in simple contour integration

**DOI:** 10.1101/350082

**Authors:** Yanli Zhou, Luigi Acerbi, Wei Ji Ma

## Abstract

Perceptual organization is the process of grouping scene elements into whole entities. A classic example is contour integration, in which separate line segments are perceived as continuous contours. Uncertainty in such grouping arises from scene ambiguity and sensory noise. Some classic Gestalt principles of contour integration, and more broadly, of perceptual organization, have been re-framed in terms of Bayesian inference, whereby the observer computes the probability that the whole entity is present. Previous studies that proposed a Bayesian interpretation of perceptual organization, however, have ignored sensory uncertainty, despite the fact that accounting for the current level of perceptual uncertainty is one the main signatures of Bayesian decision making. Crucially, trial-by-trial manipulation of sensory uncertainty is a key test to whether humans perform near-optimal Bayesian inference in contour integration, as opposed to using some manifestly non-Bayesian heuristic. We distinguish between these hypotheses in a simplified form of contour integration, namely judging whether two line segments separated by an occluder are collinear. We manipulate sensory uncertainty by varying retinal eccentricity. A Bayes-optimal observer would take the level of sensory uncertainty into account – in a very specific way – in deciding whether a measured offset between the line segments is due to non-collinearity or to sensory noise. We find that people deviate slightly but systematically from Bayesian optimality, while still performing “probabilistic computation” in the sense that they take into account sensory uncertainty via a heuristic rule. Our work contributes to an understanding of the role of sensory uncertainty in higher-order perception.

**Author summary:** Our percept of the world is governed not only by the sensory information we have access to, but also by the way we interpret this information. When presented with a visual scene, our visual system undergoes a process of grouping visual elements together to form coherent entities so that we can interpret the scene more readily and meaningfully. For example, when looking at a pile of autumn leaves, one can still perceive and identify a whole leaf even when it is partially covered by another leaf. While Gestalt psychologists have long described perceptual organization with a set of qualitative laws, recent studies offered a statistically-optimal – Bayesian, in statistical jargon – interpretation of this process, whereby the observer chooses the scene configuration with the highest probability given the available sensory inputs. However, these studies drew their conclusions without considering a key actor in this kind of statistically-optimal computations, that is the role of sensory uncertainty. One can easily imagine that our decision on whether two contours belong to the same leaf or different leaves is likely going to change when we move from viewing the pile of leaves at a great distance (high sensory uncertainty), to viewing very closely (low sensory uncertainty). Our study examines whether and how people incorporate uncertainty into contour integration, an elementary form of perceptual organization, by varying sensory uncertainty from trial to trial in a simple contour integration task. We found that people indeed take into account sensory uncertainty, however in a way that subtly deviates from optimal behavior.

## Introduction

Perceptual organization is the process whereby the brain integrates primitive elements of a visual scene into whole entities. Typically, the same scene could afford different interpretations because of ambiguity and perceptual noise. How the brain singles out one interpretation has long been described to follow a set of qualitative principles defined in Gestalt psychology. For example, contour integration, a form of perceptual organization that consists of the perceptual grouping of distinct line elements into a single continuous contour, is often described by the Gestalt principles of “good continuation” and “proximity”. These principles state that humans extrapolate reasonable object boundaries by grouping local contours consistent with a smooth global structure [1].

While Gestalt principles represent a useful catalogue of well-established perceptual phenomena, they lack a theoretical basis, cannot make quantitative predictions, and are agnostic with respect to uncertainty arising from sensory noise. This not only limits understanding at the psychological level, it is also problematic within a broader agenda of quantitatively linking neural activity in different brain areas to behavior. For example, neural investigations of the perception of illusory contours, a phenomenon in contour integration in which the observer perceives object contours when they are not physically present, have largely remained at a qualitative level. An alternative approach that does not suffer from these shortcomings uses the framework of Bayesian inference, whereby the observer computes the probabilities of possible world states given sensory observations using Bayes’ rule [2]. In the realm of perceptual organization, Bayesian models stipulate that the observer computes the probabilities of different hypotheses about which elements belong to the same object (e.g., [3–6]). For the example of contour integration, such hypotheses would be that line elements belong to the same contour and that they belong to different contours.

A fully Bayesian approach to contour integration would provide a normative way for dealing both with high-level uncertainty arising from ambiguity in the latent structure of the scene, and with low-level (sensory) uncertainty arising from noise in measuring primitive elements of the scene. Crucially, however, previous studies in perceptual organization, and more specifically contour integration, have looked at the statistics of the environment [5] but have not examined whether the decision rule adapts flexibly as a function of *sensory* uncertainty. Such adaptation is a form of *probabilistic computation* and, while not unique, is one of the basic signatures of Bayesian inference in perception [7]. This question is fundamental to understanding whether, how, and to which extent the brain represents and computes with probability distributions [8]. A trial-by-trial manipulation of sensory uncertainty is an effective test of probabilistic computation, because otherwise Bayesian inference would be indistinguishable from an observer using an inflexible, uncertainty-independent mapping [9, 10]. While the variation of sensory reliability is not the only possible form of uncertainty manipulation, it has been a successful approach for studying probabilistic computation in low-level perception, such as in multisensory cue combination [11, 12] and in integration of sensory measurements with prior expectations [13, 14]. Moreover, tasks with varying sensory uncertainty have yielded insights into the neural representation of uncertainty [15, 16].

In the current study, we investigate the effect of varying sensory uncertainty on an atomic form of contour integration. Specifically, we manipulate sensory uncertainty unpredictably on a trial-to-trial basis by changing stimulus retinal eccentricity in a simple collinearity judgment task. Our experimental manipulation allows for a stronger test of the hypothesis that perceptual grouping is a form of Bayesian inference, at least for the elementary case of collinearity judgment of two line segments. However, looking for a qualitative empirical signature of Bayesian computations, such as an effect of sensory uncertainty, is not enough because many distinct decision strategies might produce similar behaviors [17, 18]. Proper quantitative comparison of Bayesian observer models against plausible alternatives is critical in establishing the theoretical standing of the Bayesian approach [19–21]. For example, as an alternative to performing the complex, hierarchical computations characteristic of optimal Bayesian inference, the brain might draw instead on simpler non-Bayesian decision rules and non-probabilistic heuristics [22, 23] such as grouping scene elements based on some simple, learned rule. In contour integration, such a simple rule may dictate that line elements belong to the same contour if they are close enough in space and orientation, independently of other properties of the scene. Therefore, here we rigorously compare the Bayesian strategy, and sub-optimal variants thereof, against alternative and markedly non-Bayesian decision rules, both probabilistic and non-probabilistic. While we find compelling evidence of probabilistic computation, a probabilistic, non-Bayesian heuristic model outperforms the Bayes-optimal model, suggesting a form of sub-optimality in the decision-making process. Our study paves the way for a combined understanding of how different sources of uncertainty affect contour integration, and offers the opportunity for rigorous Bayesian modeling to be extended to more complex forms of perceptual organization.

## Results

Subjects (*n* = 8) performed a *collinearity judgment* task (Fig 1A). On each trial, the participant was presented with a vertical occluder and the stimulus consisted of two horizontal lines of equal length on each side of the occluder. At stimulus offset, the participant reported whether the two lines were collinear or not via a single key press. To avoid the learning of a fixed mapping, we withheld correctness feedback. In different blocks in the same sessions, participants also completed a *height judgment task* (Fig 1B), with the purpose of providing us with an independent estimate of the participants’ sensory noise. In both tasks, sensory uncertainty was manipulated by varying retinal eccentricity on a trial to trial basis (Fig 1D). We investigated whether people took into account their sensory noise *σ_x_*(*y*), which varied with eccentricity level *y*, when deciding about collinearity.

**Fig 1.**
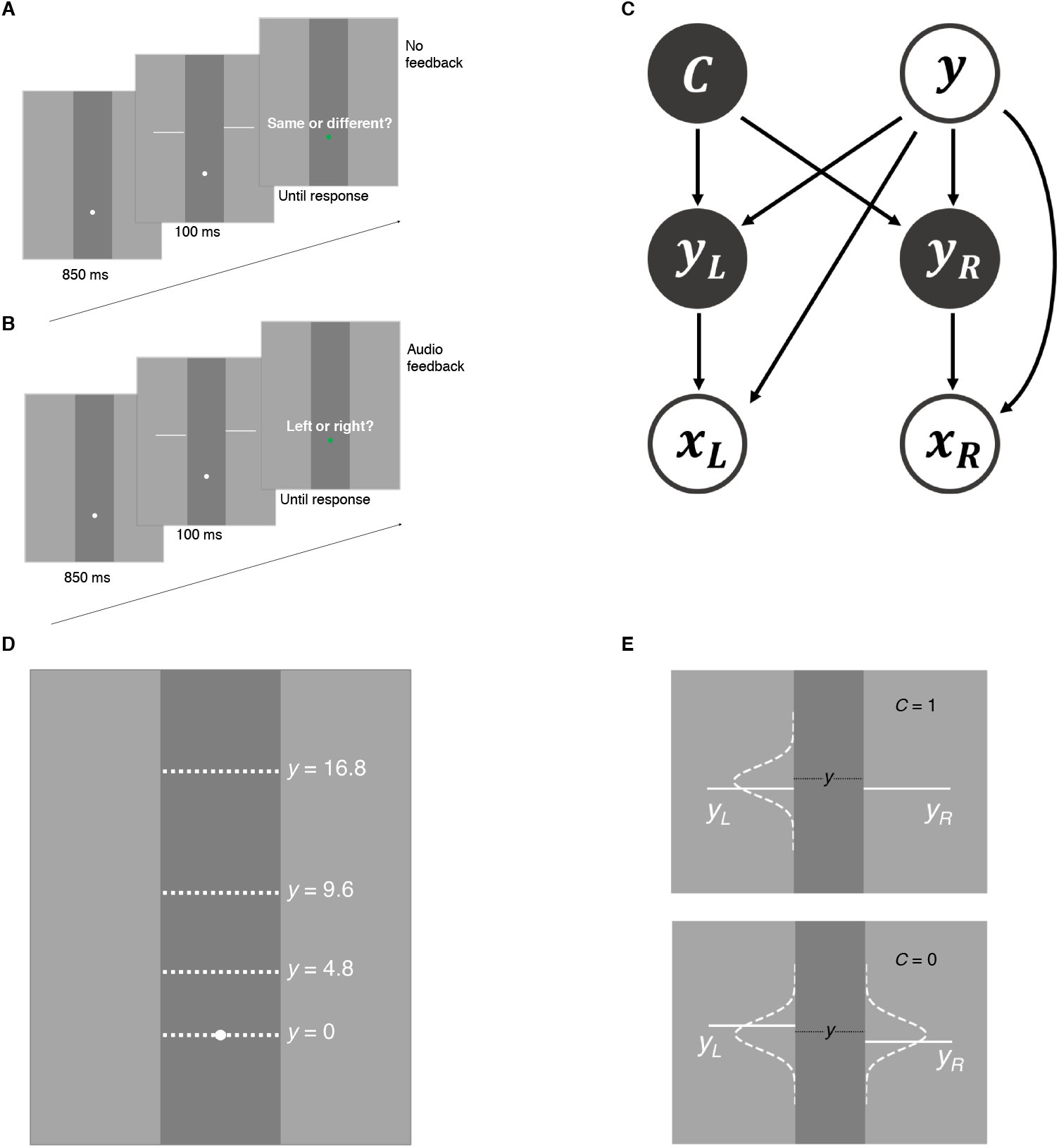
Tasks and generative model. A: Collinearity judgment task. After stimulus offset, participants reported if the line segments belonged to the same line or different lines. B: Height judgment task. Participants reported whether the left line segment was higher or the right line segment was higher. C: Generative model of the collinearity judgment task. Trial type *C* = 1 when the two lines segments are collinear, and *C* = 0 when line segments are non-collinear. On a given trial, the stimulus pair *y_L_, y_R_* randomly appeared around one of four eccentricity levels (*y* = 0, 4.8, 9.6, 16.8), measured by degrees of visual angle (dva). For all models, the observer’s measurements *x_L_, x_R_* are assumed to follow a Gaussian distribution centered on the true stimulus *y_L_, y_R_*, respectively, with standard deviation *σ_x_*(*y*) dependent on eccentricity level *y*. D: Possible eccentricity levels (in dva). E: Stimulus distribution for collinearity judgment task. When *C* = 1, the vertical position of the left line segment *y_L_* is drawn from a Gaussian distribution centered at *y* with fixed standard deviation *σ_y_*; the vertical position of the right segment *y_R_* is then set equal to *y_L_*. When *C* = 0, *y_L_* and *y_R_* are independently drawn from the same Gaussian.

We found a main effect of vertical offset on the proportion of collinearity reports (two-way repeated-measures ANOVA with Greenhouse-Geisser correction; *F*_(3.69,114)_ = 101, *ϵ* = 0.461, *p* < 0.001, 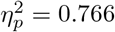) and a main effect of eccentricity (*F*_(2.38,169)_ = 51.2, *ϵ* = 0.794, *p* < 0.001, 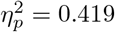), suggesting that the experimental manipulations were effective (Fig 2A,B). We also found a significant interaction between offset and eccentricity (*F*_(4.38,30.7)_ = 7.88, *ϵ* = 0.183, *p* < 0.001, 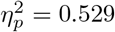), which is evident in the psychometric curves across subjects (Fig 2C).

**Fig 2.**
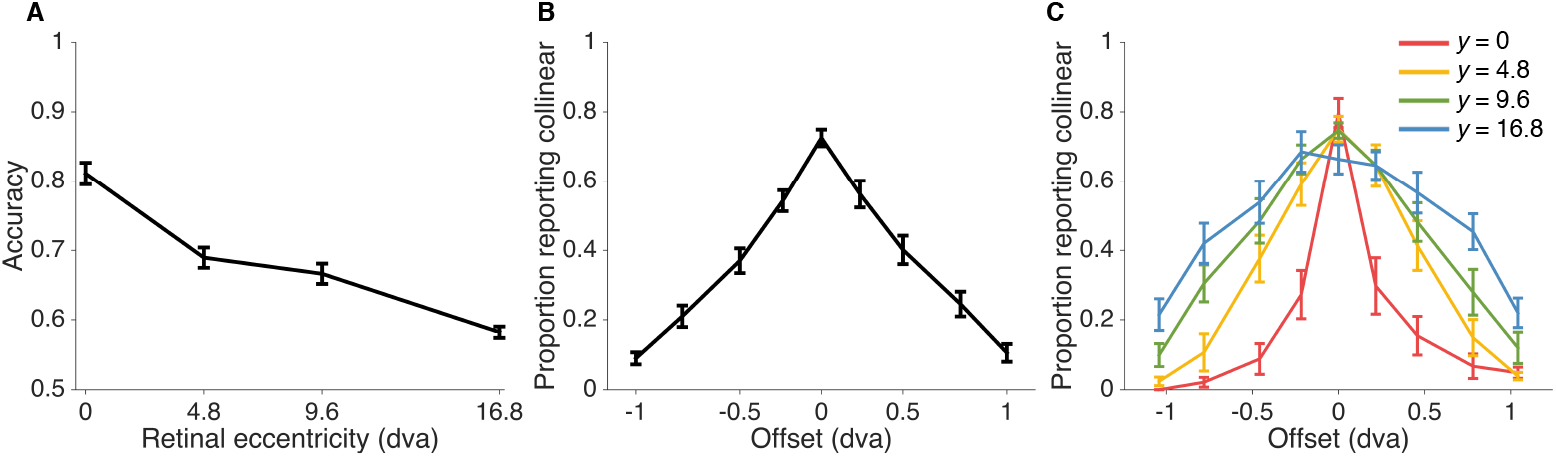
Collinearity judgement task data. A: Accuracy as a function of retinal eccentricity level (chance probability = 0.5). B: Proportion of reporting “collinear” as a function of vertical offset between the two line segments. C: Proportion of reporting “collinear” as a function of vertical offset of the two line segments at each eccentricity level. Error bars indicate Mean ± 1 SEM across 8 subjects.

We did not find significant effects of learning across sessions (see S1 Appendix), so in our analyses for each subject we pooled data from all sessions.

### Models

We describe here three main observer models which correspond to different assumptions with respect to when the observer reports “collinear”, that is three different forms of decision boundaries (Fig 3).

**Fig 3.**
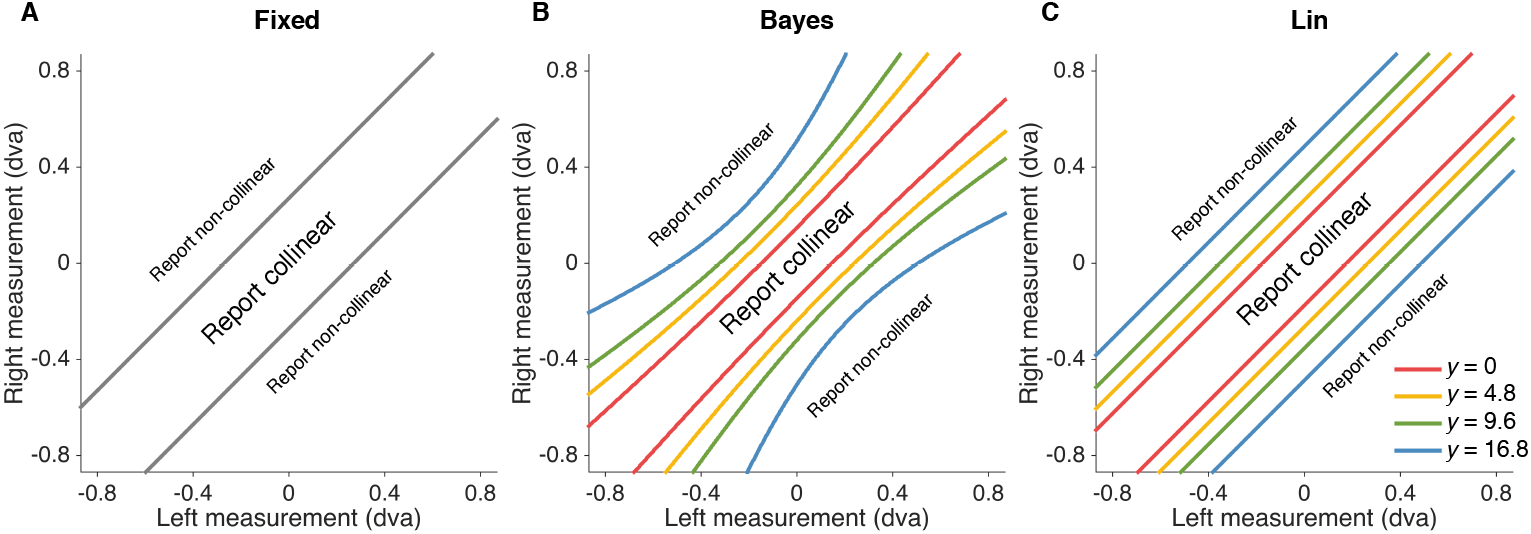
Decision boundaries for fixed-criterion (Fixed), Bayesian (Bayes) and linear heuristic (Lin) models (left to right). The probability of reporting “collinear” given stimulus and eccentricity condition is equal to the probability that the observer’s measurements of vertical positions of left and right line segments fall within the boundary defined by the model.

We first consider the behavior of a Bayesian observer (“Bayes”) who utilizes the probability distributions defined in the generative model of the task (Fig 1C) to make decisions that maximize the probability of being correct, given the available sensory measurements. In particular, the Bayesian observer accounts for uncertainty when deciding whether a measured offset between the line segments is due to non-collinearity or to sensory noise by choosing the category (*C* = 1 or *C* = 0) with the highest posterior probability *p*(*C|x_L_, x_R_*), where *x_L_, x_R_* are measurements of the two line segments on a particular trial. This strategy translates into reporting “collinear” when *x_L_*, *x_R_* fall within the Bayesian decision boundary, which is a function of (a) both measurements – not simply their difference –, (b) sensory noise (that is, eccentricity) in the trial, (c) the belief about the offset distribution width *σ_y_*, and (d) the prior belief about the proportion of collinear trials *p*(*C* = 1) (Fig 3B). Note that a strictly Bayes-optimal observer would have a prior that matches the experimental distribution, *p*(*C* = 1) = 0.5. Here we relaxed the assumption and allowed *p*(*C* = 1) to be a free parameter. The basic Bayesian model assumes that the observer knows the width of the offset distribution (fixed throughout the experiment) and the noise level associated with the current trial; see Model variants for a relaxation of these assumptions.

To investigate whether people apply instead a learned stimulus mapping that is uncertainty independent, we tested a fixed-criterion model (“Fixed”) [24] in which the observer responds that two line segments are collinear whenever the measured offset |*x_L_* − *x_R_*| is less than a fixed distance *κ* (a free parameter of the model). This corresponds to an eccentricity-invariant decision boundary (Fig 3A).

Finally, we also considered an instance of probabilistic, non-Bayesian computation via a heuristic model (“Lin”) in which the observer takes stimulus uncertainty into account in a simple, linear way: the observer responds “collinear” whenever the measured offset |*x_L_* − *x_R_*| is less than an uncertainty-dependent criterion,

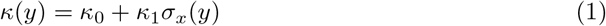

where *κ*_0_ and *κ*_1_ are free parameters of the model (Fig 3C). While the Lin model takes uncertainty into account, and thus it is “probabilistic”, it is formally non-Bayesian because it does not use knowledge of the statistics of the task to compute a posterior over latent variables [7] (see also Discussion).

A detailed mathematical description of each model is reported in S1 Appendix.

### Model comparison

To fully account for parameter uncertainty, we used Markov Chain Monte Carlo (MCMC) to sample the posterior distributions of the parameters for each model and individual subject. To estimate goodness of fit (that is, predictive accuracy) while taking into account model complexity, we compared models using the leave-one-out cross-validation score (LOO), estimated on a subject-by-subject basis directly from the MCMC posterior samples via Pareto smoothed importance sampling [25] (see Methods). Higher LOO scores correspond to better predictive accuracy and, thus, better models.

We found that the fixed-criterion model fits the worst (LOO_Bayes_ − LOO_Fixed_ = 25.6 ± 13.6, LOO_Lin_ − LOO_Fixed_ = 69.3 ± 16.5; Mean ± SEM across subjects), while also yielding the poorest qualitative fits to the behavioral data (Fig 4A). This result suggests that participants used not only their measurements but also sensory uncertainty from trial to trial, thus providing first evidence for probabilistic computation in collinearity judgment. Moreover, we find that the linear heuristic model performs better than the Bayesian model (LOO_Lin_ − LOO_Bayes_ = 43.7 ± 13.3), suggestive of a suboptimal way of taking uncertainty into account.

**Fig 4.**
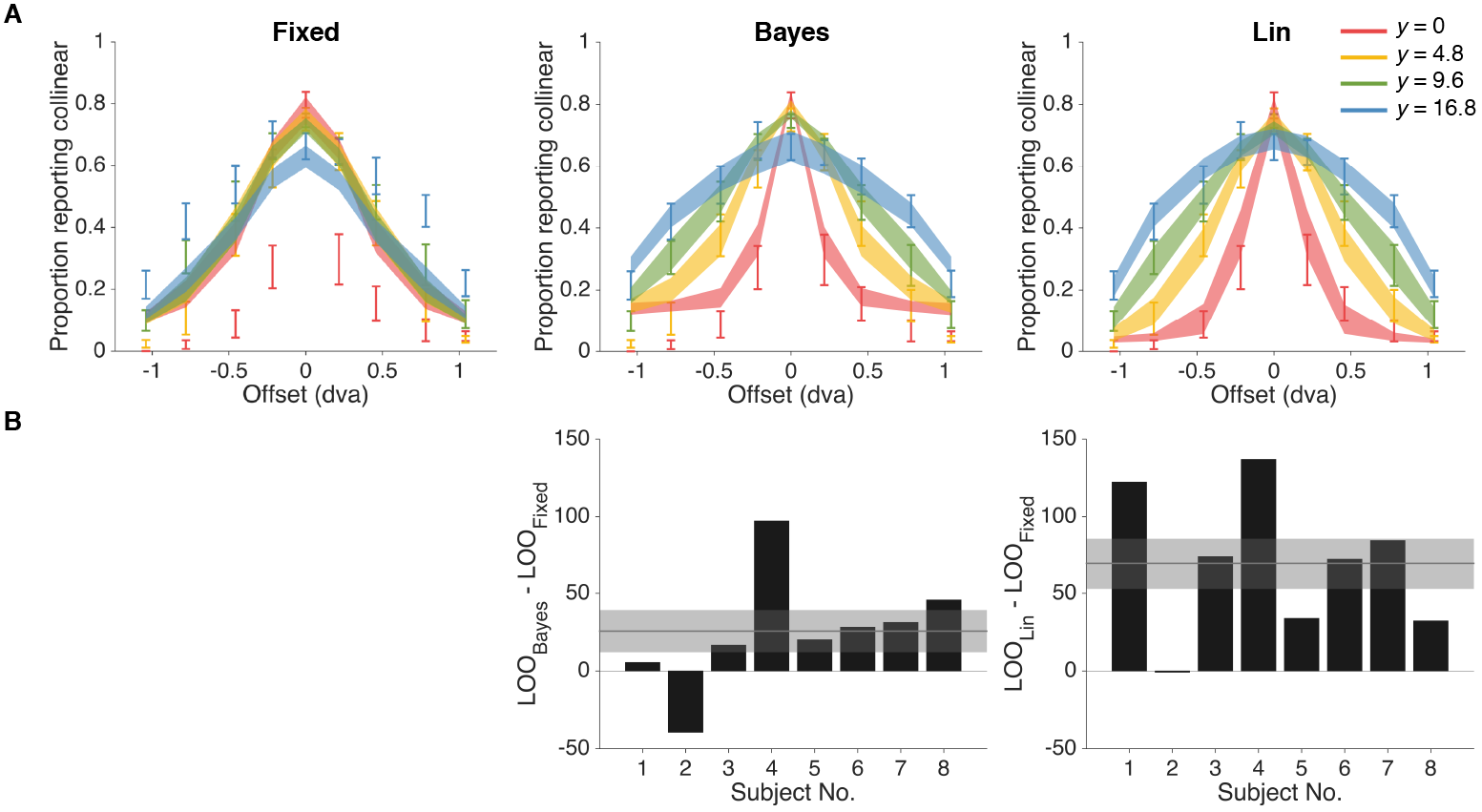
Model fits and model comparison for fixed-criterion (Fixed), Bayesian (Bayes) and linear heuristic (Lin) models (from left to right). A: Model fits to proportion of responding collinear as a function of vertical offset of the two line segments. Error bars indicate Mean ± 1 SEM across subjects. Shaded regions indicates Mean ± 1 SEM of fits for each model, with each model on a separate column. B: Model comparison via leave-one-out cross-validation score (LOO). Bars indicate individual subjects’ LOO scores for every model, relative to the fixed-criterion model. A positive value indicates that the model in the corresponding column had a better LOO score than the fixed-criterion model. Shaded regions indicate Mean ± 1 SEM in LOO differences across subjects. The Lin model won the model comparison, whereas Fixed was the worst model.

To allow for model heterogeneity across subjects, we also combined model evidence from different subjects using a hierarchical Bayesian approach that treats the model as a random variable to accommodate between-subject random effects [26]. This method allowed us to compute the expected posterior frequency for each model, that is the probability that a randomly chosen subject belongs to a particular model in the comparison. This analysis confirmed our previous model comparison ordering, with the Fixed model having the lowest expected frequency (0.11 ± 0.09), Bayes the second highest (0.18 ± 0.11) and Lin by far the highest (0.71 ± 0.13). We also calculated the protected exceedance probability [27], that is the probability that a particular model is the most frequent model in the set, above and beyond chance. We found consistent results – namely the Fixed model has the lowest protected exceedance probability (0.048), followed by Bayes (0.062), and Lin (0.89).

### Validation of noise parameters

In all analyses so far, the observer’s sensory noise levels at each eccentricity level *σ_x_*(*y*) were individually fitted as free parameters (four noise parameters, one per eccentricity level). To obtain an independent estimate of the subjects’ noise, and thus verify if the noise parameters estimated from the collinearity task data truly capture subjects’ sensory noise, we introduced in the same sessions an independent Vernier discrimination task (height judgment task) [28, 29]. In this task, participants judged whether the right line segment was displaced above or below the left line segment (Fig 1B and Fig 5A). Importantly, the observer’s optimal decision rule in this task is based solely on the sign of the observer’s measured offset between the line segments, and does not depend on the magnitude of sensory noise (that is, respond “right segment higher” whenever *x_R_* > *x_L_*). Moreover, trials in this task matched the stimulus statistics used in non-collinear trials of the collinearity judgment task. Therefore, the height judgment task afforded an independent set of estimates of subjects’ noise levels.

**Fig 5.**
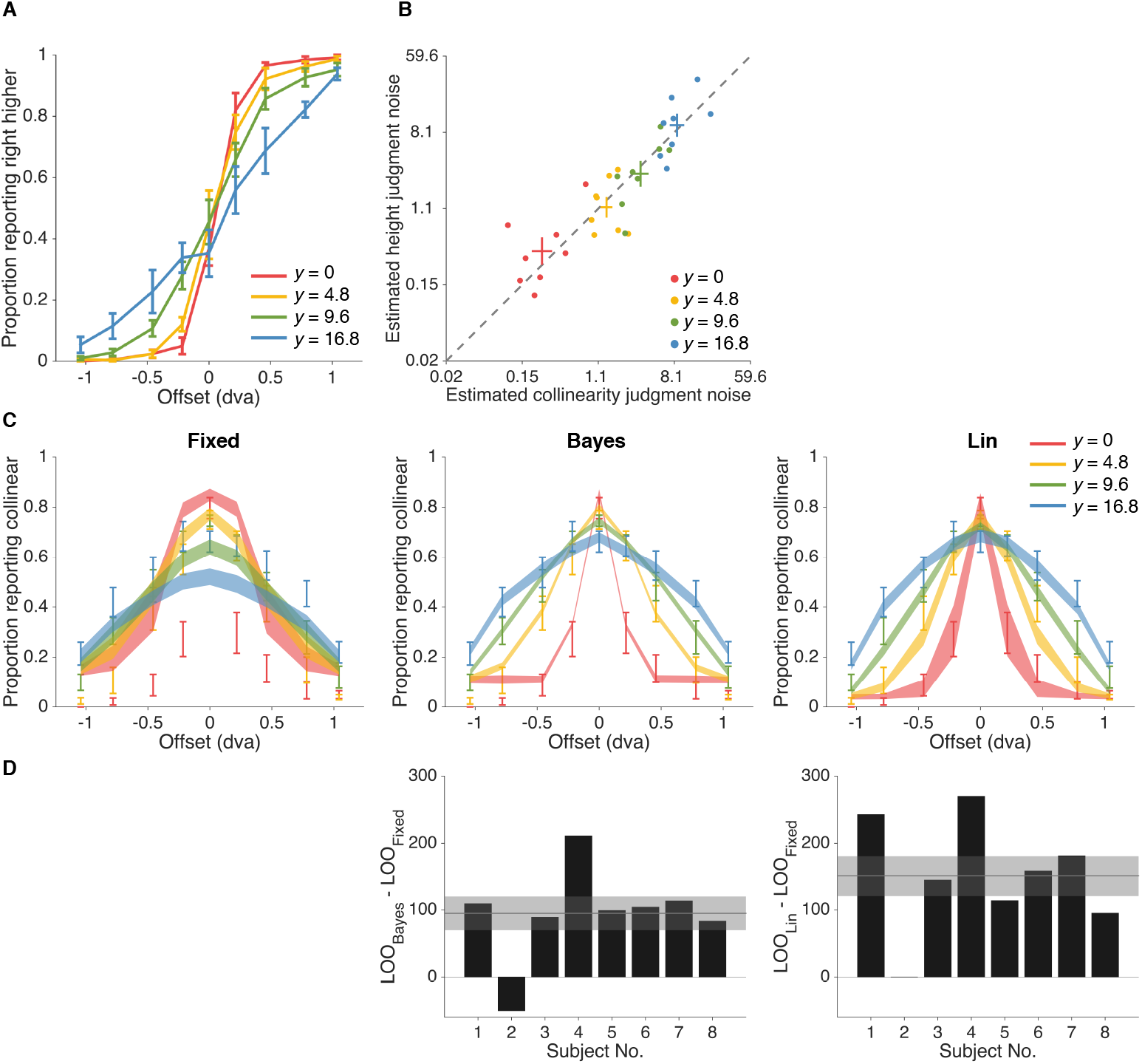
Height judgment task results. A: Height judgment task data. Proportion of reporting “right line segment higher” is plotted as a function of vertical offset between line segments. Error bars indicate Mean ± 1 SEM across subjects. B: Noise parameters estimated from the best-fitting model, linear heuristic (Lin), on collinearity judgment task vs. noise parameters estimated from the height judgment task, in dva. Each dot corresponds to a subject’s estimated noise parameters (posterior means) for a given eccentricity level. C: Models’ fits to collinearity judgment task data when noise parameters estimated from the height judgment task were imported into the models. Shaded regions indicate Mean ± 1 SEM of fits. See Fig 4A for comparison. D: Model comparison on collinearity judgment task data via LOO, constrained by importing noise parameters from the height judgment task. Results are consistent with the model comparison ordering we found in the original unconstrained fits, with free noise parameters (see Fig 4B for comparison).

Repeated-measures ANOVA indicated a main effect of the vertical offset between the two line segments on the proportion of reports “right higher” (*F*_(2.58,80.1)_ = 320, *ϵ* = 0.323, *p* < 0.001, 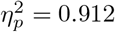), no main effect of eccentricity (*F*_(1.99,141)_ = 0.300, *ϵ* = 0.662, *p* = 0.740, 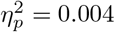), and an interaction between eccentricity and offset (*F*_(4.67,32.7)_ = 8.75, *ϵ* = 0.195, *p* < 0.001, 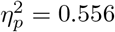). These findings confirm that, as expected, participants in the height judgement task took into account the offset, and their performance was also affected simultaneously by offset and eccentricity (that is, sensory noise).

We found that sensory noise parameters estimated from the best model (Lin) in the collinearity task were well correlated – across subjects and eccentricities – with those estimated from the height judgment task (*r* = 0.88) (Fig 5B), indicating that the model is correctly capturing subjects’ noise characteristics in collinearity judgment.

We next examined whether the model comparison between Bayes, Fixed, and Lin could be constrained using the parameter estimates obtained from the height judgment task, and whether such a constrained comparison would alter our findings. For each subject and each eccentricity level, we imported the posterior mean of each noise parameter of that subject at that eccentricity level, as estimated from the height judgment task, into the model for the collinearity task. This left the Bayes, Fixed, and Lin models with only 2, 2, and 3 free parameters, respectively, which we estimated via MCMC as previously described. The fits of the constrained models were comparable to those of their unconstrained counterparts (compare Fig 5C to 4A). The quantitative comparison of the constrained models was also consistent with that of the unconstrained models (compare Fig 5D to 4B): LOO_Bayes_ − LOO_Fixed_ = 94.7 ± 25.1, LOO_Lin_ − LOO_Fixed_ = 150.6 ± 30.2. Overall, this analysis shows that our models correctly captured subjects’ noise features, and that our conclusions are not merely due to excessive flexibility of our models, as we obtain the same results with models with very few free parameters.

As a further sanity check, we repeated the analysis in this section using maximum-a-posteriori (MAP) estimates for the noise parameters imported from the height judgment task (instead of the posterior means), finding quantitatively similar results (correlation between collinearity task and height judgement task parameters: *r* = 0.87; LOO_Bayes_ − LOO_Fixed_ = 93.5 ± 26.7; LOO_Lin_ − LOO_Fixed_ = 142.4 ± 28.3).

### Suboptimality analysis

In the previous sections we have found that the Lin model wins the model comparison against the Bayesian model, suggestive of suboptimal behavior among participants. Here we closely examine the degree of suboptimality in terms of the loss of accuracy in the collinearity task with respect to Bayes-optimal behavior.

In order to assess the accuracy that an observer with a given set of noise parameters could achieve, had they performed Bayes-optimally, we proceeded as follows. For each subject, we generated a simulated dataset from the Bayesian model using the maximum-a-posteriori noise parameters *σ_x_*(*y*) estimated from both the collinearity judgment task and the height judgment task. We used both estimates to ensure that our results did not depend on a specific way of estimating noise parameters. For this analysis, we assumed optimal parameters, that is *p*_common_ = 0.5 and no lapse (*λ* = 0).

We found a significant difference between observed accuracy and estimated optimal accuracy based on collinearity judgment noise, as shown in Fig 6 (two-way prepeated-measures ANOVA with Greenhouse-Geisser correction; *F*_(1.00,7.00)_ = 37.8, *ϵ* = 1.00, *p* < 0.001, 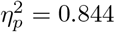). There is a significant main effect of eccentricity (*F*_(2.03,14.2)_ = 128, *ϵ* = 0.675, *p* < 0.001, 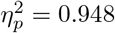), which is expected from the experimental manipulations. We also found no significant interaction between optimality condition and eccentricity (*F*_(2.22,15.5)_ = 2.31, *ϵ* = 0.738, *p* = 0.106, 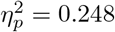). Analogously, for height judgement noise parameters, there is also a significant difference between observed accuracy and estimated optimal accuracy (*F*_(1.00,7.00)_ = 7.45, *ϵ* = 1.00, *p* = 0.029, 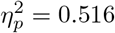), a significant main effect of eccentricity (*F*_(1.85,13.0)_ = 54.4, *ϵ* = 0.618, *p* < 0.001, 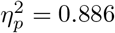), and no significant interaction between optimality condition and eccentricity (*F*_(2.16,15.1)_ = 3.46, *ϵ* = 0.720, *p* = 0.055, 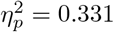). These results confirm the results of the model comparison in that there is a statistically significant difference between our subjects’ performance and optimal behavior.

**Fig 6.**
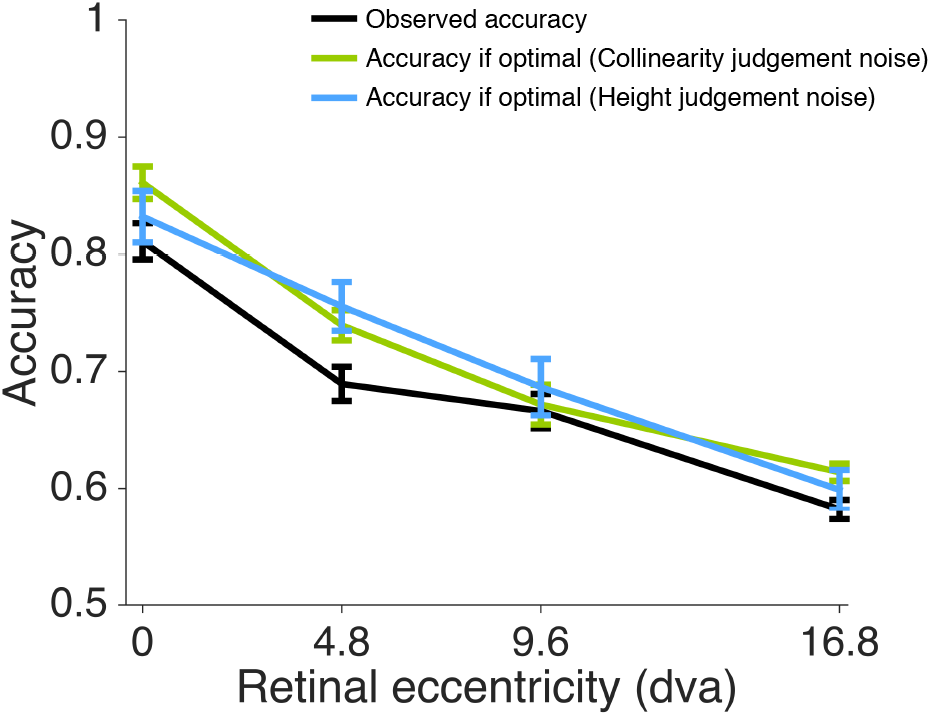
Suboptimality analysis. Black line: Observed accuracy across four eccentricity levels (chance probability = 0.5). Error bars indicate Mean ± 1 SEM across subjects. Green line: Estimated accuracy if subjects perform Bayes-optimally, with noise parameters obtained via the collinearity judgement task. Blue line: Estimated accuracy with noise parameters obtained via the height judgment task. Performance was slightly suboptimal across participants.

However, a *statistically* significant difference does not necessarily imply a substantial difference in terms of performance, as previous studies have shown that participants can ±be “optimally lazy” by deviating from optimal performance in a way that has minimal impact on overall expected score in a task [30]. We quantified our subjects’ performance in terms of *efficiency*, that is the proportion of correct responses with respect to optimal behavior. Our subjects exhibited an overall efficiency of 0.953 ± 0.007 (based on collinearity judgment noise), or 0.959 ± 0.015 (based on height judgement noise), which suggests that our subjects were only slightly suboptimal (see Discussion).

### Model variants

We consider here several alternative observer models that relax some key assumptions we made when constructing our main observers, to verify whether our findings still hold.

#### Trial dependency of sensory uncertainty

We tested for potential influence of stimulus uncertainty from previous trials (“History” model) on the response of the current trial. Specifically, for the History model we extended the formula of the decision boundary of the Lin model to be a linear function of the noise parameters of the current trial, as before, plus the noise associated with up to four previous trials, that is *σ_x_*(*y_t_*), *σ_x_*(*y*_*t*−1_), *σ_x_*(*y*_*t*−2_), *σ_x_*(*y*_*t*−3_), *σ_x_*(*y*_*t*−4_), respectively, each one with a separate weight.

We found no evidence of trial dependency based on sensory uncertainty, for the History model fits about as well or even slightly worse than Lin (LOO_History_ − LOO_Lin_ = −2.4 ± 0.24). In particular, we also found that the maximum-a-posteriori weights associated with *σ_x_*(*y_t−_*_1_) to *σ_x_*(*y_t−_*_4_) were all not significantly different from zero across participants (respectively, *t*_(7)_ = 1.45, *p* = 0.19; *t*_(7)_ = 0.0754, *p* = 0.94; *t*_(7)_ = −1.18, *p* = 0.28; *t*_(7)_ = −1.27, *p* = 0.24). These results show that sensory uncertainty from previous trials had no effect on the observers’ decision in the current trial.

#### Mismatch of noise parameters

So far, we have assumed that observers utilize directly their noise parameters *σ_x_*(*y*) when computing the decision rule. Here we propose a variant of the Bayesian model, “Mismatch”, in which the observer instead uses a set of *assumed* noise parameters that may deviate from the true standard deviations of their measurement distributions [31]. This model is identical to the Bayesian model except that all four *σ_x_*(*y*) are substituted with *σ_x,_*_assumed_(*y*), the assumed noise parameters, in the calculation of the decision variables. To limit model complexity, we chose for the assumed noise parameters a parametric form which is a linear function of the true noise parameters *σ_x_*(*y*). To avoid issues of lack of parameter identifiability [17], for the Mismatch model we also fixed *p*_common_ = 0.5. Thus, the Mismatch model has the same number of free parameters as Lin, and one more than Bayes.

After relaxing the Bayesian model to allow for assumed noise parameters, we found that the Mismatch model fits better than the original Bayes model 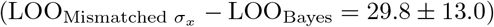, and, thus, better than the Fixed model as well, which was already the worst in the comparison 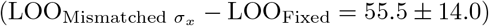. However, we found that the Lin model is still the best-fitting model 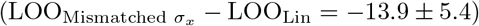. All combined, these results suggest that a degree of suboptimality in the observers might have arisen from a lack of knowledge of their own noise characteristics [31], but such mismatch is not enough to entirely explain the observed pattern of behavior.

#### Mismatch of stimulus distribution width

We consider another relaxation of the Bayesian observer model, whereby the subject computes with an incorrect stimulus distribution width *σ*_*y*,assumed_, which governs the offset distribution (Fig 1E) and is fixed throughout the experiment. In the main Bayesian model, the observer is assumed to have learned the true value of *σ_y_* from training trials presented during the experiment. However, it is possible that a lack of attention during training or a lack of training itself could lead to a mismatched estimation of *σ_y_*, which then influences the calculation of the decision boundaries.

The results of this Width-mismatched model indicate that incorporating the incorrect assumption of stimulus distribution width also effectively improves the performance of the Bayesian model 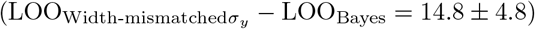 while still losing to the Lin model 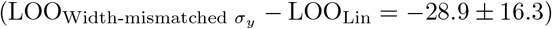. We see that the deviation from optimality observed in the data might also be partially attributed to the lack of familiarity with the stimulus distribution, yet still not enough to better account for the data than our current best model, namely the Lin model.

#### Bayesian observer with decision noise

In the basic Bayesian model, we model the inference stage as exact. That is, we assume there is no noise or imperfection in the mapping from the observer’s internal representation to the decision. We now consider a formulation of the Bayesian model in which the observer performs Bayesian inference with decision noise [32]. Specifically, we model decision noise *σ_d_* as a Gaussian noise on the decision variable [33–36]:

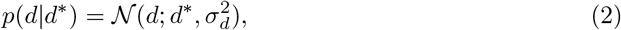

where *d*\* is the original decision variable in the basic Bayesian model (see S1 Appendix).

Consistent with previous analyses, we see that this new relaxation of the original Bayesian model with decision noise (Bayes+DN) better accounts for the behavioral data than the basic Bayesian observer (LOO_Bayes+DN_ − LOO_Bayes_ = 25.4 ± 15.2), but is still not able to outperform the Lin model (LOO_Bayes+DN_ − LOO_Lin_ = 18.3 ± 8.2).

Despite having tried different forms of relaxation of the basic Bayesian model, we still find the Lin model to be our best performing model. Finally, taking our approach one step further, we also tested a suboptimal Bayesian model in which we allowed for both stimulus distribution width mismatch and decision noise. Having 8 free parameters, one more than the Lin model, the hybrid model can be expected to be a very flexible model. However, while once again improving on performance within the Bayesian formulation 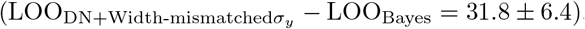, the hybrid model is still could not surpass the Lin model 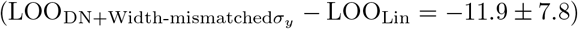.

#### Nonparametric examination

In the Lin model (and variants thereof), so far we assumed a linear parametric relationship between the decision boundary and the noise level *σ_x_*(*y*), as per Eq 1.

Here we loosened this constraint and fitted the decision boundary for each eccentricity level as an individual free parameter. Due to its flexible nature, we consider this “Nonparametric” model merely as a descriptive model, which we expect to describe the data very well. We use the Nonparametric model as a means to provide an upper-bound on the LOO score for each individual, so as to have an absolute metric to evaluate the performances of other models (in a spirit similarly to estimating the entropy of the data, that is an estimate of the intrinsic variability of the data which represents an upper bound on the predictive performance of any model [37, 38]). As expected, given the large amount of flexibility, the Nonparametric model fits better than Lin (LOO_Nonparametric_ − LOO_Lin_ = 14.6 ± 6.5), but we note that the difference in LOO is substantially less than the difference between Lin and Bayes (43.7 ± 13.3), or Lin and Fixed (69.3 ± 16.5), suggesting that Lin is already capturing subjects’ behavior quite well, close to a full nonparametric description of the data.

We can also use the Nonparametric model to examine how close the parametric estimates of decision boundary from Lin, our best model so far, are to those obtained nonparametrically. We observed that the average decision boundary across 8 subjects, as a function of eccentricity, was consistent with the average nonparametric estimates of the decision boundary at every eccentricity level (Fig 7A,B). This agreement means that the decision boundaries adopted by observers in the task were, indeed, approximately linear in the sensory noise associated with each eccentrity level, as assumed by the linear heuristic model (Eq 1), thus validating our modeling choice.

**Fig 7.**
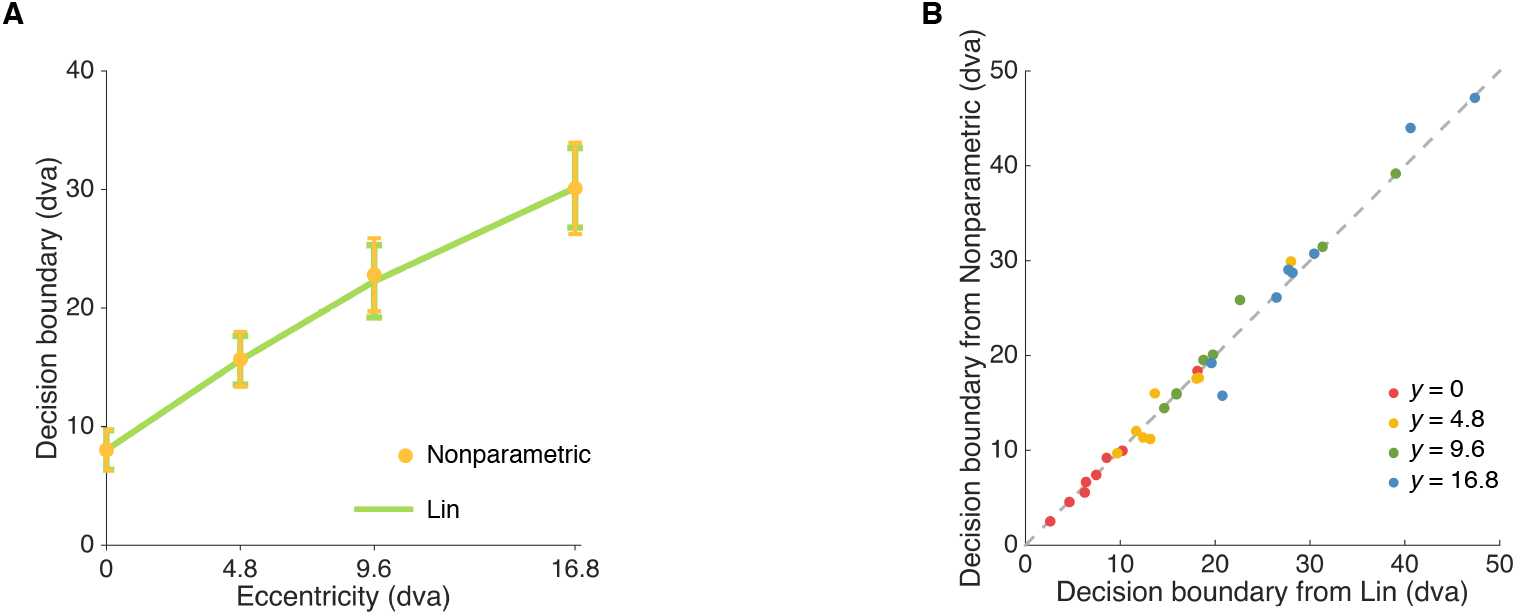
Nonparametric model. A: Decision boundary estimates of linear heuristic model (Lin) vs. decision boundary estimates of the Nonparametric model at different eccentricity levels (Mean ± 1 SEM). B: Decision boundary at every eccentricity level fitted non-parametrically vs. Decision boundary at every eccentricity level fitted from the Lin model. Even when allowed to vary freely (“non-parametrically”), the decision boundaries are approximately linear in the sensory noise associated with each eccentricity level (and, incidentally, approximately linear in the eccentricity level itself), as per the Lin model.

## Discussion

To study how people group together local elements to form a continuous contour, we designed a behavioral experiment in which participants were asked to judge whether two line segments partially occluded belonged to the same line. Using computational observer models to describe the obtained data, we found that people utilize sensory uncertainty when making collinearity judgements, however in a slightly suboptimal way.

Crucially, our results are robust to changes in model assumptions, such as noise model mismatch, history effects, and different decision boundaries, and we independently validated our parameter estimates in a different task. With trial-by-trial manipulation of eccentricity in a collinearity judgment task, our study presents a rigorous examination of the role of sensory uncertainty for probabilistic computations in contour integration.

### Contour integration as elementary perceptual organization

The present study is linked to the broader effort to study hierarchical Bayesian inference in perception, whereby the observer is required to marginalize over stimulus values (here, line offset) to build a posterior over latent, discrete causal scenarios (here, same line of different lines). Such framework was adopted and tested in a variety of domains such as cue combination [39], change detection [33], perception of sameness [40], and causal inference [41]. In particular, our models share the same formal structure of models of causal inference in multisensory perception [41, 42]. In such tasks, the observer receives sensory measurements of possibly discrepant cues from distinct sensory modalities (e.g., vision and hearing), and has to infer whether the cues originated from the same source (*C* = 1) or from different sources (*C* = 0) – leading to, respectively, cue integration and cue segregation. Previous work has shown that Bayesian causal inference models provide a good qualitative description of human performance in multisensory perception with discrepant cues, but quantitative comparison hints at deviations from exact Bayesian behavior [38], not unlike what we find here. Our study differs from previous work in that here we focus on an atomic form of perceptual organization.

Our approach is related to the work by Stevenson and Körding [43] in the field of depth perception, which also provides an example of uncertainty manipulation used to probe the basis of elementary perceptual organization. In their task, observers inferred whether a stereographically displayed disk was occluded or not, and utilized that information about the configuration of the visual scene to match the depth of another disk. Importantly, the experiment included two different sensory uncertainty conditions. Biases in depth estimation showed by subjects in the two conditions were well explained by a Bayesian model that inferred the relative probability of the two causal scenarios. Crucially, however, Stevenson and Körding did not test alternative models that could also account for the data.

While in our study we closely examined the effect of varying sensory uncertainty, our task did not strictly introduce ambiguity, a integral element of Gestalt perception. Ambiguity translates to overlapping stimulus distributions, and ambiguous trials are only found in the collinear category of our task. With the presence of ambiguity, an observer will not be able to achieve perfect performance even when sensory noise is completely absent. Shapes defined by illusory contours such as variants of the Kanizsa triangle were previously used to study representations of illusory contours in the cortical areas of the brain in functional imaging [44, 45], rendering them potential candidates for stimuli that can incorporate both ambiguity and sensory uncertainty.

Nevertheless, by studying the role of sensory uncertainty alone, our study presents a more careful account of Bayesian inference in contour integration, and potentially in perceptual organization. In particular, we compared the Bayesian observer model against other observer models that each describe an alternative plausible decision strategy. We were able to distinguish a fixed stimulus mapping that mimics Bayesian inference from probabilistic computation, which requires the observer to flexibly adjust their decision boundary according to sensory uncertainty. Despite evidence for probabilistic computations, we found that data was better explained by a non-Bayesian heuristic model (but see below for further discussion).

Using simple contour integration as a starting point, our study provides detailed examinations of the Bayesian observer model in comparison to other models, paving the way for the applications of similarly rigorous analyses on more complex and naturalistic forms of perceptual organization. Our paradigm can be modified to examine the effects of varying occluder width, varying contour angles, thickness and curvature to simulate the complexity of real-world contour statistics [5]; and to test whether humans utilize the task statistics when computing the posterior probability of different explanations. Even in naturalistic perceptual organization tasks, Bayesian strategies are still favorable for their inherent ability to control for model complexity, due to the process of marginalizing (i.e., integrating) over latent variables, which tends to favor the simplest hypothesis (property often referred to as ‘Bayesian Occam’s razor’ [46]). Notably, the model comparison pipeline laid out in the current study can be easily extended to be applied to these tasks.

### Heuristic and imperfect strategies in probabilistic contour integration

Our findings align with a variety of results that show that, in many perceptual decision making tasks, humans perform moderately to reasonably well, but not quite in a Bayes-optimal manner [47]. In particular, extended and rigorous model comparison often reveals that human behavior can be better explained by observer models representing strategies or heuristics which are not exactly Bayesian [31, 36, 38, 48–50] (but see [37] for an example where the Bayes-optimal model wins against a wide array of contenders).

However, the difference between a probabilistic, non-Bayesian heuristic (which may be interpreted as “approximating Bayesian inference”) and an imperfect, noisy or mismatched Bayesian model is arguably moot. When people deviate from ‘correct’ Bayesian behavior, it can be very hard – and it might even become impossible in principle – to fully distinguish between (a) mismatched Bayesian computations (i.e., due to ‘model mismatch’ between the observer’s beliefs and the true statistics of the task); (b) approximate Bayesian computations (due to implementations of approximate inference in the brain, necessary to deal with any complex models of the environment, and with limited resources [51]); (c) the adoption of some other complex heuristics with certain Bayesian-looking signatures (e.g., taking uncertainty into account). Even when the behavior *looks* fairly Bayesian, as for example according to a probabilistic strategy that takes uncertainty into account in an approximately correct way, we may be unable to distinguish “truly” Bayesian computations (that explicitly account for uncertainty according to Bayes’ rule) from arbitrarily complex non-Bayesian strategies that use complex heuristics and properties of the stimulus to estimate the current trial uncertainty. The problem is ill-posed in psychophysics as there is always some property of the stimulus which correlates with trial uncertainty, and we cannot know the exact computations being performed in the brain until we open the black box and look at the actual neural implementation.

The empirical question that we *can* address via psychophysics is how flexible the subjects’ strategies are in accounting for uncertainty, and how close they end up being to the Bayesian strategy, independently of the source of uncertainty, and even when multiple sources of uncertainty are combined [52]. Moreover, we can use psychophysical data to investigate why (and how) subjects deviate from the optimal performance, and how the brain *learns* to process sensory information so as to perform a given task, which is an avenue for future work.

In our case, a possible explanation for subjects’ heuristic strategy, which differed slightly but systematically from optimal performance, might be that they had received insufficient training. While we found no evidence of learning across sessions, it is possible that participants would have learnt to perform optimally had they received correctness feedback on the task, possibly with greater incentives to motivate their learning. The main purpose of our experiment was to explore the role of sensory uncertainty – thus, we limited the amount of training trials with performance feedback on purpose, to prevent the possible learning of a fixed mapping of stimulus to collinearity condition that is independent of sensory uncertainty. The tradeoff between providing sufficient training trials and avoiding learning of fixed mapping makes it difficult to test behaviorally the hypothesis that sub-optimality stems from insufficient training.

A possible alternative avenue for exploring the effect of task learning could be through training an artificial neural network on the same psychophysical task, and examining how performance evolves as a function of training epochs [53], and whether this mimics human behavior. For example, a hierarchical, probabilistic and stochastic neural network such as Deep Boltzmann Machine is a desirable candidate as it can learn to generate sensory data in an unsupervised fashion, a procedure that provides a plausible account for visual cortical processing [54, 55]. Notably, such stochastic hierarchical generative model was used to show that visual numerosity – a higher-order feature – can be invariantly encoded in the deepest hidden layer of the neural network [56], and could analogously give rise to illusory contours neurons as found in monkeys [54].

In conclusion, our finding that elementary contour integration is probabilistic – albeit slightly suboptimal – leads naturally to a fundamental open question in neuroscience, that is whether and how the visual system performs (or approximates) probabilistic inference in the presence of complex, naturalistic stimuli. There is a trade-off between stimulus complexity and modeling tractability in that we experimenters do not normally have access to the generative model of a complex visual scene, preventing the deployment of powerful statistical tools from ideal-observer analysis such as those used in the current work. However, for example, a recent theoretical paper introduced a flexible, parametric model of overlapping and occluded geometric shapes that resemble the pattern of a bed of leaves (“dead leaves” [57]). Our rigorous model comparison approach, combined with such complex psychophysical stimuli, provides a viable direction for future studies interested in further exploring the probabilistic nature of perceptual organization.

## Methods

### Ethics statement

The Institutional Review Board at New York University approved the experimental procedures (protocol #IRB-FY2016-599: “Visual perception, attention, and memory”) and all subjects gave written informed consent.

### Subjects

8 subjects (6 female), aged 20-30, participated in the experiment. Subjects received $10 for each of four 1-hour sessions, plus a completion bonus of $10.

### Apparatus and stimuli

The stimuli were shown on a 60 Hz 9.7-inch 2048-by-1536 pixel display. The display (LG LP097QX1-SPA2) was the same as that used in the 2013 iPad Air (Apple). The screen was secured to an arm with height adjusted to each subject’s eye level. A chin rest was horizontally aligned with the center of the screen. The distance between the eyes and the display was 27.5 cm. To minimize potential biases caused by external visual cues, we added a large black panel surrounding the display. The display was connected to a Windows desktop PC using the Psychophysics Toolbox extensions [58, 59] for MATLAB (MathWorks, Natick, MA).

On each trial, a dark gray occluder (23 cd*/*m^2^) with a width of 5.6 degrees of visual angle (dva) was displayed against a light gray background (50 cd*/*m^2^). A white (159 cd*/*m^2^) fixation dot 0.24 dva in diameter was shown in the lower central part of the occluder; this dot corresponded to a retinal eccentricity of 0 dva. The stimuli consisted of two horizontal white line segments on both sides of the occluder. The line segments were all 5.6 dva in width and 0.16 dva in height. The vertical “base position” *y* of a pair of line segments had one of four levels of retinal eccentricity (0, 4.8, 9.6, and 16.8 dva).

### Trial procedure

Subjects completed two tasks, which we call *collinearity judgment* task and *height judgment* task. On each trial in the collinearity judgment task (Fig 1A), the occluder and fixation dot were displayed for 850 ms, followed by the stimulus for 100 ms. On a “non-collinear” trial, the vertical positions of the two line segments were independently drawn from a normal distribution centered at one of the four “base” eccentricity levels (0, 4.8, 9.6, or 16.8 dva), with a standard deviation of 0.48 dva (Fig 1E); on a “collinear” trial, we drew the vertical position of the line segment on one side and matched the line segment on the other side. In each session, 50% of the trials were “collinear” and 50% were “non-collinear”, randomly interleaved. At stimulus offset, the fixation dot turned green to prompt the subject to indicate whether the two line segments were collinear. The participant pressed one of 8 keys, corresponding to 8 choice-confidence combinations, ranging from high-confident collinear to high-confident non-collinear. Response time was not constrained. No performance feedback was given at the end of the trial.

Height judgment task trials followed the same procedure (Fig 1B), except that the subject was asked to report which of the two line segments was highest (“left” or “right”). We generated the line segments in the same fashion as in the “non-collinear” condition of the collinearity judgment task. Audio feedback was given after each response to indicate whether the choice was correct.

For the analyses described in this paper, we only considered choice data (“collinear/non-collinear”, “left/right”), leaving analysis of confidence reports to future work.

### Experiment procedure

During each session, subjects completed one height judgment task block, followed by three collinearity judgment task blocks, and finished with another height judgment task block. Each height judgment task block consisted of 60 trials, and each collinearity judgment task block consisted of 200 trials.

A demonstration of the experimental procedure was given to each subject at the beginning of the first session. Participants were informed that there were an equal number of left/right trials in the height judgment task as well as an equal number of collinear/non-collinear trials in the collinearity judgment task. To familiarize subjects with the stimulus distribution and to check for understanding of the tasks, participants completed 16 practice trials at the beginning of each session. Stimulus presentation time was longer on practice trials (500 ms), and audio correctness feedback was given at the end of each practice trial. We did not analyze the responses on the practice trials.

### Data analysis

In order to visualize psychometric curves with enough trials, we binned the offset values between the left and right line segments into the following intervals: (−∞, −3.31], (−3.31, −2.08], (−2.08, −1.17], (−1.17, −0.38], (−0.38, −0.38], (0.38, 1.17], (1.17, 2.08], (2.08, 3.31], (3.31, ∞), in dva. These values were chosen to include a comparable number of trials per interval, based on the quantiles of the Gaussian distribution of the offset used in the experiment. For the collinearity judgment task, we computed the proportion of trials in which subjects reported “collinear” at each offset bin and retinal eccentricity level. For the height judgment task, we computed the proportion of trials in which subjects reported “right higher” at each offset bin and retinal eccentricity level.

Repeated-measures ANOVA with offset bin and eccentricity level as within-subjects factors were performed separately on the proportion of reporting “collinear” in the collinearity judgment task and the proportion of reporting “right higher” in the height judgment task. We applied Greenhouse-Geisser correction of the degrees of freedom in order to account for deviations from sphericity [60], and report effect sizes as partial eta squared, denoted with 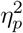.

For all analyses the criterion for statistical significance was *p* < .05, and we report uncorrected *p*-values. Unless specified otherwise, summary statistics are reported in the text as mean ± SEM between subjects. Note that we used the summary statistics described in this section only for visualization and to perform simple descriptive statistics; all models were fitted to raw trial data as described next.

### Model fitting

For each model and subject, the noise parameters 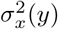 for *y* = 0, 4.8, 9.6 and 16.8 dva were fitted as individual parameters.

We calculated the log likelihood of each individual dataset for a given model with parameter vector ***θ*** by summing the log probability of trial *i* over all N trials,

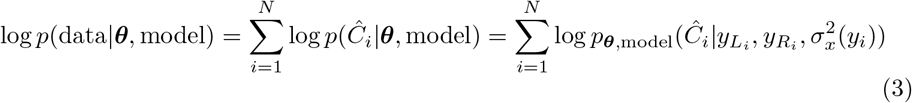

where the response probability 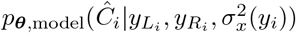 is defined in S1 Appendix.

We fitted the models by drawing samples from the unnormalized log posterior distribution of the parameters *p*(***θ***|data) using Markov Chain Monte Carlo (parallel slice sampling [38, 61]) for each subject. The posterior distribution of the parameters is proportional to the sum of data likelihood (Eq 4) and a factorized prior over each parameter *j*,

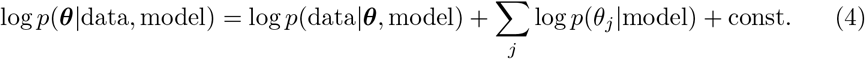

We used log-transformed coordinates for scale parameters (e.g., noise), and for all parameters we assumed a uniform non-informative prior (uniform in log space for scale parameters) [62], within reasonably large bounds. Three parallel chains were ran with starting point set at maximum likelihood point estimates of the parameters, evaluated with Bayesian Adaptive Direct Search [63], to ensure that the chains were initialized within a high posterior density region.

After running all chains, we computed Gelman and Rubin’s potential scale reduction statistic *R* for all parameters to check for convergence [64]. An *R* value that diverges from 1 indicates convergence problems, whereas a value close to 1 suggests convergence of the chains. The average difference between *R* value and 1 across all parameters, subjects and models is 1.16 × 10^−4^, and all *R* values fall within (0.99, 1.003], suggesting good convergence. To verify compatibility between different runs, we also visually inspected the posteriors from different chains. We merged samples from all chains for each model and subject in further analyses.

To visualize model fits (or posterior predictions) in Fig 4 and 5, we computed the posterior mean model prediction for each subject based on 60 independent samples from the posterior (equally spaced in the sampled chains). We then plotted average and standard deviation across subjects.

### Model comparison

To estimate the predictive accuracy of each model while taking into account model complexity, we performed Bayesian leave-one-out (LOO) cross-validation. Bayesian LOO cross-validation computes the posterior of the parameters given *N* − 1 trials (the training set), and evaluates the (log) expected likelihood of the left-out trial (the test set); this process is repeated until all trials have been iterated through, yielding the leave-one-out score

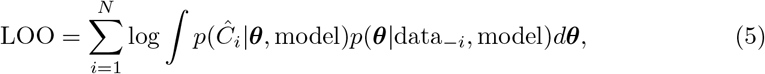

where *p*(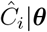, model) is the likelihood of the *i*-th trial (see Eq 3), and *p*(***θ***|data_−*i*_, model) is the posterior over ***θ*** given all trials except the *i*-th one. For most models, evaluating Eq 5 naively is impractical due to the cost of obtaining the leave-one-out posteriors via *N* distinct MCMC runs. However, noting that all posteriors differ from the full posterior by only one data point, one could approximate the leave-one-out posteriors via *importance sampling*, reweighting the full posterior obtained in a single MCMC run. Still, a direct approach of importance sampling can be unstable, since the full posterior is typically narrower than the leave-one-out posteriors. Pareto-smoothed importance sampling (PSIS) is a recent but already widely used technique to stabilize the importance weights [25]. Thus, Eq 5 is approximated as

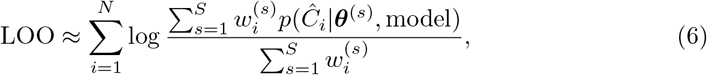

where *θ*^(*s*)^ is the *s*-th parameter sample from the posterior, and 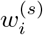 are the Pareto-smoothed importance weights associated to the *i*-th trial and *s*-th sample (out of *S*); see [25] for details and [38] for an application with discussion of other model selection metrics.

We also conducted model comparison by computing a simpler metric, namely the Akaike information criterion (AIC) using log likelihood values obtained via maximum likelihood estimation for each subject and each model. We found a perfect rank correlation between AIC scores and LOO scores (*ρ* = 1), which is expected in the limit of infinite trials, for AIC and LOO cross-validation are asymptotically equivalent [65].

### Data and code

Subjects’ data sets and the code used to perform our model fitting and comparison analyses are available with instructions at the following link: https://github.com/yanlizhou/collinearity

## Acknowledgments

We thank Andra Mihali, Will Adler, Maija Honig and Zahy Bnaya for useful discussions of earlier versions of this manuscript. This work has utilized the NYU IT High Performance Computing resources and services.

## S1 Appendix

Supplemental methods. Analysis of learning; model specification; model recovery analysis; posterior distributions of model parameters.

## Notes

#### Summary of Updates

1. We analyzed and compared in the main text several additional "suboptimal" Bayesian observer models, in particular: A. a model with mismatched belief about the width of the stimulus (offset) distribution (Bayes Width-mismatched); B. model with noise on the decision variable (Bayes+DN); C. a model that has both sources of suboptimality introduced above (Bayes Width-mismatched + DN). While these models generally perform better than the basic Bayesian model, they still do not explain the data better than the “heuristic” Lin model. Thus, our main conclusions are unaffected. 2. We performed the "noise transfer" analysis again by using posterior mean estimates for the parameters, in addition to maximum-a-posteriori (MAP) estimates, finding quantitatively similar results. We report both in the Results, as a way to confirm the robustness of the analysis. 3. To better reflect the scope of our work, we clarified that the focus of the manuscript is more on an elementary form of "contour integration" than on "perceptual organization" as a whole. We modified accordingly the Title, Abstract, Author Summary, Introduction and Discussion. 4. In the Methods section we largely extended the explanation of Pareto-Smoothed Importance Sampling leave-one-out cross-validation (PSIS-LOO), which is the technique we used to compute our main model comparison metric (LOO). 5. Still, we verified that our results do not rely on us using a specific model comparison metric, by checking that findings are the same under another metric (AIC). Relatedly, we also performed our model recovery analysis using AIC. 6. We added to Appendix S1 a new figure that shows the posterior distributions of the model parameters obtained via MCMC for the three main models (Bayes, Fixed, and Lin), for a representative subject. 7. We extended the Discussion to address in detail some of the points made by the reviewers (e.g., distinguishability of Bayesian vs. heuristic models; discussing related work on occlusion) 8. We added (and fixed) several references; we fixed typos; we clarified a few ambiguous sentences; and implemented a number of edits and suggestions.

https://github.com/yanlizhou/collinearity

