## Supplementary material for "The role of sensory uncertainty in simple contour integration": S1 - Appendix

### Appendix S1
### Supplemental methods

Yanli Zhou[☯]*, Luigi Acerbi[☯], Wei Ji Ma

☯ These authors contributed equally to this work.
* Contact:

**3 Model recovery analysis** **4**

**4 Posterior distributions of model parameters** **4**

#### 1 Analysis of learning

We found no main effect of learning across sessions on participants' accuracy in the collinearity task, as shown in Fig S1 (two-way repeated-measures ANOVA with Greenhouse-Geisser correction; $F_{(2.49,17.4)} = 0.859$, $\epsilon = 0.828$, $p = 0.462$, $\eta_p^2 = 0.109$). There is a significant main effect of eccentricity ($F_{(2.28,16.0)} = 62.4$, $\epsilon = 0.761$, $p < 0.001$, $\eta_p^2 = 0.899$), which is expected from the experimental manipulations. We also found no significant interaction between session and eccentricity ($F_{(3.44,24.1)} = 0.624$, $\epsilon = 0.382$, $p = 0.627$, $\eta_p^2 = 0.082$). These analyses suggest that participants quickly learnt the task and their performance was stationary across sessions.

#### 2 Model specification

##### 2.1 Bayesian model

In each trial, an observer utilizes the noisy measurements $x_L, x_R$ of the true stimuli $y_L, y_R$ to produce an estimate of the collinearity state $\hat{C}$. Under the Bayesian model, the observer accounts for sensory

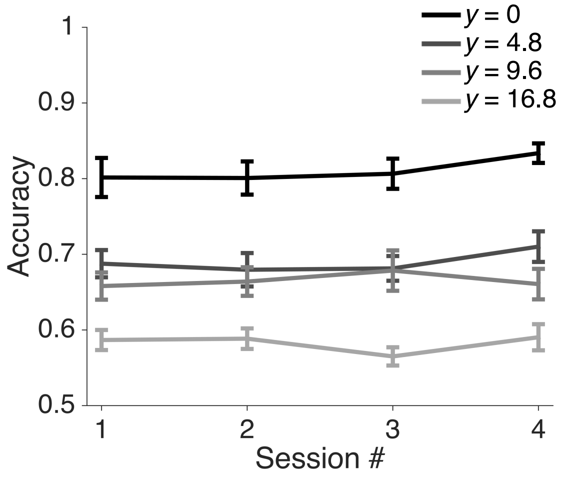

**Figure S1. Analysis of learning across sessions.** Accuracy across four sessions (chance probability = 0.5). Error bars indicate Mean ± 1 SEM across subjects, with retinal eccentricity level plotted as separate lines. We found no significant change in performance across sessions.

uncertainty when deciding whether a measured offset between the line segments is due to non-collinearity or to sensory noise by computing the log posterior ratio $d(x_L, x_R)$ (Eq S1) of the two competing hypotheses so that the probability of answering correctly is maximized,

$$d(x_L, x_R) = \log \frac{p(C=1|x_L, x_R)}{p(C=0|x_L, x_R)} = \log \frac{p(x_L, x_R|C=1)}{p(x_L, x_R|C=0)} + \log \frac{p(C=1)}{p(C=0)}. \tag{S1}$$

Having no direct knowledge of the true vertical positions, the observer marginalizes over $y_L$ and $y_R$. This gives rise to the following expression for $d(x_L, x_R)$,

$$d(x_L, x_R) = \log \frac{\mathcal{N}(x_L; x_R, 2\sigma_x^2)\mathcal{N}(\frac{x_L+x_R}{2}; y, \frac{\sigma_x^2}{2} + \sigma_y^2)}{\mathcal{N}\left(x_L; y, \sigma_x^2 + \sigma_y^2\right)\mathcal{N}(x_R; y, \sigma_x^2 + \sigma_y^2)} + \log \frac{p(C=1)}{1 - p(C=1)}, \tag{S2}$$

where $\mathcal{N}\left(x|\mu, \sigma^2\right)$ is the probability density of a normal distribution with mean $\mu$ and variance $\sigma^2$, and we left out the $y$-dependence of $\sigma_x^2(y)$ for notational simplicity.

The Bayesian decision rule is to report "collinear" when $d(x_L, x_R) > 0$,

$$\hat{C} = \begin{cases} 1 & \text{if } d > 0 \\ 0 & \text{if } d < 0 \end{cases} \tag{S3}$$

The boundary equality $d(x_\mathrm{L}, x_\mathrm{R}) = 0$ defines a curve in $(x_\mathrm{L}, x_\mathrm{R})$-space, which we visualize in Fig 3B of the main text for several values of $\sigma_x^2$.

#### 2.2 Fixed-criterion model

We also tested a non-Bayesian model in which the observer applies a fixed, uncertainty-independent decision boundary $\kappa$ (Fig 3A in the main text). The estimated collinearity state is determined as follows,

$$\hat{C} = \begin{cases} 1 & \text{if } |\Delta x| > \kappa \\ 0 & \text{if } |\Delta x| < \kappa \end{cases} \tag{S4}$$

This is equivalent to using a decision variable $d(x_L, x_R) = |x_L - x_R| - \kappa$. This model describes the strategy in which the observer uses only the measurements of the line segments, and reports collinearity if the measured offset is within a fixed threshold $\kappa$.

#### 2.3 Linear heuristic model

Our third main model is a non-Bayesian probabilistic model in which the decision boundary is represented by a linear function of sensory noise (Fig 3C in the main text),

$$\hat{C} = \begin{cases} 1 & \text{if } |\Delta x| > \kappa_0 + \kappa_1 \sigma_x \\ 0 & \text{if } |\Delta x| < \kappa_0 + \kappa_1 \sigma_x \end{cases} \tag{S5}$$

This is equivalent to using a decision variable $d(x_L, x_R) = |x_L - x_R| - k_0 - k_1 \sigma_x$. This observer accounts for uncertainty but not in the optimal way.

#### 2.4 Response probabilities

The probability of reporting collinearity given stimuli $y_L$ and $y_R$ and eccentricity level $y$ is then equal to the probability that the measurements fall within the boundary defined by the model $M$,

$$\begin{aligned} p_M\left(\hat{C} = 1 \left| y_L, y_R, \sigma_x^2\right.\right) &= \iint p(\hat{C} = 1 | x_L, x_R, M) p(x_L | y_L) p(x_R | y_R) dx_L dx_R \\ &= \iint_{d_M(x_L, x_R) \geq 0} \mathcal{N}(x_L; y_L, \sigma_x^2) \mathcal{N}(x_R; y_R, \sigma_x^2) dx_L dx_R. \end{aligned} \tag{S6}$$

Eq S6 generally does not have an analytical solution for an arbitrary decision rule $d_M(x_L, x_R)$, in which case we estimated the response probabilities via numerical integration on a grid.

#### 2.5 Lapse rate

For all models, we fitted a lapse rate $\lambda$ for each subject. We define a lapse as a trial in which the subject randomly reports collinearity or non-collinearity, each with a probability of 0.5.

Adding the lapse rate to Eq S6 we obtain

$$p_{M,\text{lapse}}\left(\hat{C} = 1 \left| y_L, y_R, \sigma_x^2\right.\right) = (1 - \lambda) \cdot p_M\left(\hat{C} \left| y_{L_i}, y_{R_i}, \sigma_x^2\right.\right) + \frac{\lambda}{2}. \tag{S7}$$

#### 2.6 Height judgment task model

During the height judgment task, the observer reports whether the right line segment is higher than the left line segment. The observer's decision $\hat{C}$ = "right higher" or $\hat{C}$ = "left higher" depends only on the sign of the offset between measurements $\Delta x = x_R - x_L$, which in turn depends on the offset between the true vertical positions of the stimuli $\Delta y = y_R - y_L$ at the current eccentricity level. The decision rule is simply to report "right higher" when $\Delta x > 0$ and vice versa,

$$\hat{C} = \begin{cases} \text{"right higher"} & \text{if } \Delta x > 0 \\ \text{"left higher"} & \text{if } \Delta x < 0 \end{cases} \tag{S8}$$

The response probability is therefore

$$\begin{aligned} p(\hat{C} = \text{"right higher"} | \Delta y) &= \int p(\hat{C} = \text{"right higher"} | \Delta x) p(\Delta x | \Delta y) d\Delta x \\ &= \int \chi(\Delta x > 0) \mathcal{N}(\Delta x; \Delta y, 2\sigma_x^2) d\Delta x, \end{aligned} \tag{S9}$$

where $\chi(\cdot)$ is the indicator function which is equal to 1 when the argument is true, 0 otherwise.

#### 3 Model recovery analysis

To ensure our models are distinguishable, and as a further validation of our model fitting pipeline, we performed a model recovery analysis which verifies that the fitted models are matched to the data-generating model for simulated data. In this analysis, we considered our three main models: Fixed, Bayes and Lin.

We first generated 50 synthetic datasets from each of these three models using parameter vectors randomly drawn from the posterior samples of a randomly drawn subject obtained through MCMC sampling, so as to represent 'typical' observers. Following this procedure, we generated 150 datasets in total (3 generating models $\times$ 50 simulated observers). We then fitted all models to each generated dataset via maximum likelihood estimation for computational tractability. Here we used the Akaike Information Criterion (AIC) as the metric for model comparison, and computed the fraction of times that a model wins the model comparison for a given generating model (Fig S2). We did not use LOO scores for model comparison due to the impracticality of performing MCMC on all synthetic datasets and models, as we found AIC scores and LOO scores to be highly consistent for our models and datasets (see main text).

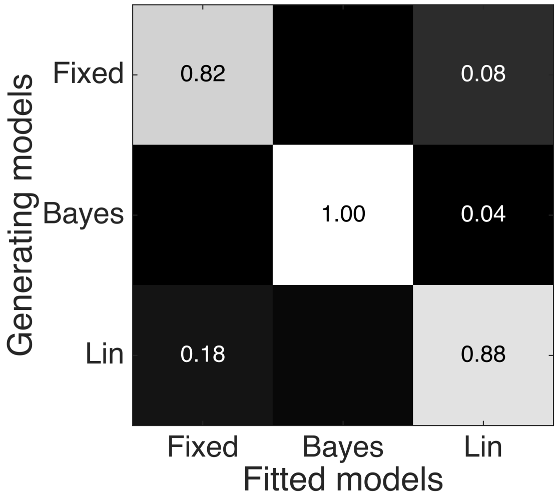

**Figure S2. Model recovery analysis.** Each square represents the fraction of datasets that were best fitted from a model (columns), for a given generating model (rows), according to the AIC score. The light shade of the diagonal squares indicates that the true generating model was, on average, the best-fitting model in all cases, leading to a successful model recovery.

On average, 90.0% of the 150 datasets were successfully recovered. This result suggests that our main models are distinguishable, and also validates our model fitting pipeline.

#### 4 Posterior distributions of model parameters

In our analyses, we first performed maximum-likelihood estimation for every model and every subject. We then followed up with MCMC sampling, using the MLE solutions as starting points for the MCMC chains. This procedure enabled us to not only obtain point estimates of model parameters for each subject but also samples from the posterior landscapes of all model parameters (examples shown below).

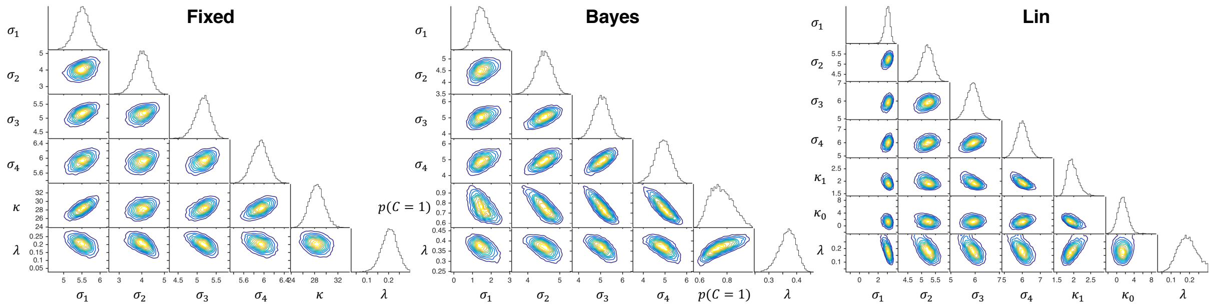

**Figure S3. Posterior distribution landscapes.** From left to right, the figure shows the posterior distributions of the model parameters obtained via MCMC for the three main models (Bayes, Fixed, and Lin, respectively), for a representative subject. In each panel, the plots represent the 1-D marginal posterior distributions of each parameter (on the diagonal), and 2-D marginal posteriors for every parameter pair (below the diagonal).
